# Modifying the trajectory of epicardial leads can substantially reduce MRI-induced RF heating in pediatric patients with a cardiac implantable electronic device

**DOI:** 10.1101/2022.07.26.501576

**Authors:** Fuchang Jiang, Bhumi Bhusal, Bach Nguyen, Michael Monge, Gregory Webster, Daniel Kim, Giorgio Bonmassar, Andrada R. Popsecu, Laleh Golestanirad

## Abstract

**Purpose:** Infants and children with congenital heart defects, inherited arrhythmia syndromes, and congenital cardiac conduction disorders often receive epicardial implantable electronic devices. Unfortunately, once an epicardial device is implanted, the patient is no longer eligible to receive MRI exams due to an elevated risk of RF heating. Here we show that a simple modification in the trajectory of epicardial leads can substantially and reliably reduce RF heating during MRI at 1.5 T, with benefits extending to abandoned leads.

**Methods:** Electromagnetic simulations were performed to assess RF heating of two common epicardial lead trajectories exhibiting different degrees of coupling with MRI incident electric fields. Experiments in anthropomorphic phantoms implanted with commercial cardiac implantable electronic devices (CIEDs) confirmed the findings.

**Results:** Simulations of an epicardial lead with a trajectory where the excess length of the lead was looped and placed on the anterior surface of the heart showed a 9-fold reduction in 0.1g-averaged SAR compared to the lead with excess length looped on the inferior surface of the heart. Repeated experiments with a commercial epicardial device confirmed the results, showing a 16-fold reduction in the average temperature rise for fully implanted systems with leads following low-SAR trajectories, and a 20-fold reduction in RF heating on an abandoned lead.

**Conclusion:** Surgical modification of epicardial lead trajectory can substantially reduce RF heating at 1.5 T, with benefits extending to abandoned leads.

## Introduction

Congenital heart defects (CHD) are the most common birth defect in the United States, affecting 1 in every 100 babies born yearly (1). Infants and children with CHD may require cardiac implantable electronic devices (CIEDs) such as pacemakers and implantable cardioverter-defibrillators, some receiving an implant within hours, or even minutes, from birth (2). The optimal approach to affixing a CIED to the heart of a young patient is to open the chest and surgically suture the cardiac lead to the myocardium (“epicardial leads”), as opposed to passing it through veins and affixing it to the inside of the heart (“endocardial leads”). Unfortunately, once epicardial leads have been implanted, patients face restrictions against MRI exams due to the elevated risk of radiofrequency (RF) heating of epicardial leads (3–5). MR-conditional CIEDs with endocardial leads have been approved by the FDA for adults, allowing patients to receive MRIs under conditions that assure safety. However, no MR-conditional system is currently available for children where epicardial leads are used more frequently than in adults (5). This leaves the most vulnerable patient population unable to receive the same standard of care as adults. In addition, children with CHD often require complex decision-making and would derive particular benefits from MRI’s sensitivity.

Advances in medical device technology have greatly mitigated the risks associated with static and gradient fields. This includes, for example, the reduction of ferromagnetic material to reduce the risk of device dislodgement due to static magnetic fields (6) and enhanced device programming to reduce the risk of cardiac stimulation from MRI-induced gradient currents (7). RF-induced heating, on the other hand, remains a safety issue. The main concern is the “antenna effect”, a phenomenon in which the electric field of the transmit RF couples with implanted leads and amplifies the specific absorption rate (SAR) of the radiofrequency energy in the tissue surrounding the lead’s tip (8, 9). The strength of this coupling depends, among other factors, on the relative orientation of the lead with respect to the transmit electric field (E). As such, the trajectory of an implanted lead substantially affects its MR-induced RF heating (10–15). This has important ramifications for patient safety as surgical guidelines are silent about how to position the excessive length of implanted leads within the body. Consequently, surgeons place the leads based on personal experience and surgical convenience, causing a substantial patient-to-patient variation in the trajectory of the lead and, by proxy, large variations in MR-induced RF heating.

Recent studies in patients with deep brain stimulation (DBS) devices have shown that modification of lead trajectory can substantially reduce MRI-induced RF heating (10,12,16). Specifically, it was shown that introducing loops in the extracranial trajectory of a DBS lead reduced local SAR at the lead tips during MRI at 1.5 T and 3 T and that the degree of the reduction in SAR depends on the location of the loop on the skull (12,16). Here we investigated if the concept of lead trajectory modification could be extended to reduce RF heating of epicardial leads in pediatric CIED patients, that is, if specific epicardial lead trajectories are more favorable in terms of MRI safety. To do this, we first performed electromagnetic simulations to identify lead trajectories that exhibit reduced coupling with the incident E at 1.5 T. We then quantified the RF power deposition in the tissue surrounding tips of leads routed along low and high coupling trajectories and verified our findings experimentally in a multimaterial anthropomorphic pediatric phantom implanted with a commercial epicardial CIED. Finally, we performed a sensitivity analysis to assess the effect of perturbations in lead trajectories and examined whether safety gains due to trajectory modification would translate to abandoned epicardial leads.

Our work is the first to suggest that an easy-to-implement surgical modifications can substantially and reliably reduce RF heating of epicardial leads in children.

### Theoretical framework

Two types of CIED leads are commonly used in clinical practice depending on the patient’s anatomy and body size: Adults and older children typically receive endocardial systems, where the leads are passed through the subclavian vein to reach the interior of the heart, with the implantable pulse generator (IPG) placed in the left or right subpectoral pocket (Figure 1 A). Infants and young children (who have small veins) mostly receive an epicardial system, which requires sternotomy to sew the cardiac lead directly to the myocardium with the IPG placed deep to the rectus abdominus muscle (Figure 1B). Because venous anatomy is relatively similar among the population, the trajectory of endocardial leads is not highly variable from patient to patient. In contrast, there is a much larger degree of freedom in placing epicardial leads, giving rise to patient-to-patient variation in epicardial lead trajectories. As the magnitude and phase of the tangential component of MRI incident electric field along a lead’s trajectory are shown to be indicators of lead’s RF heating (13,14,17–19), one would expect to see a large variation in RF heating of epicardial leads; conversely, new opportunities to increase patient safety through surgical modification of lead trajectories.

**Figure 1:**
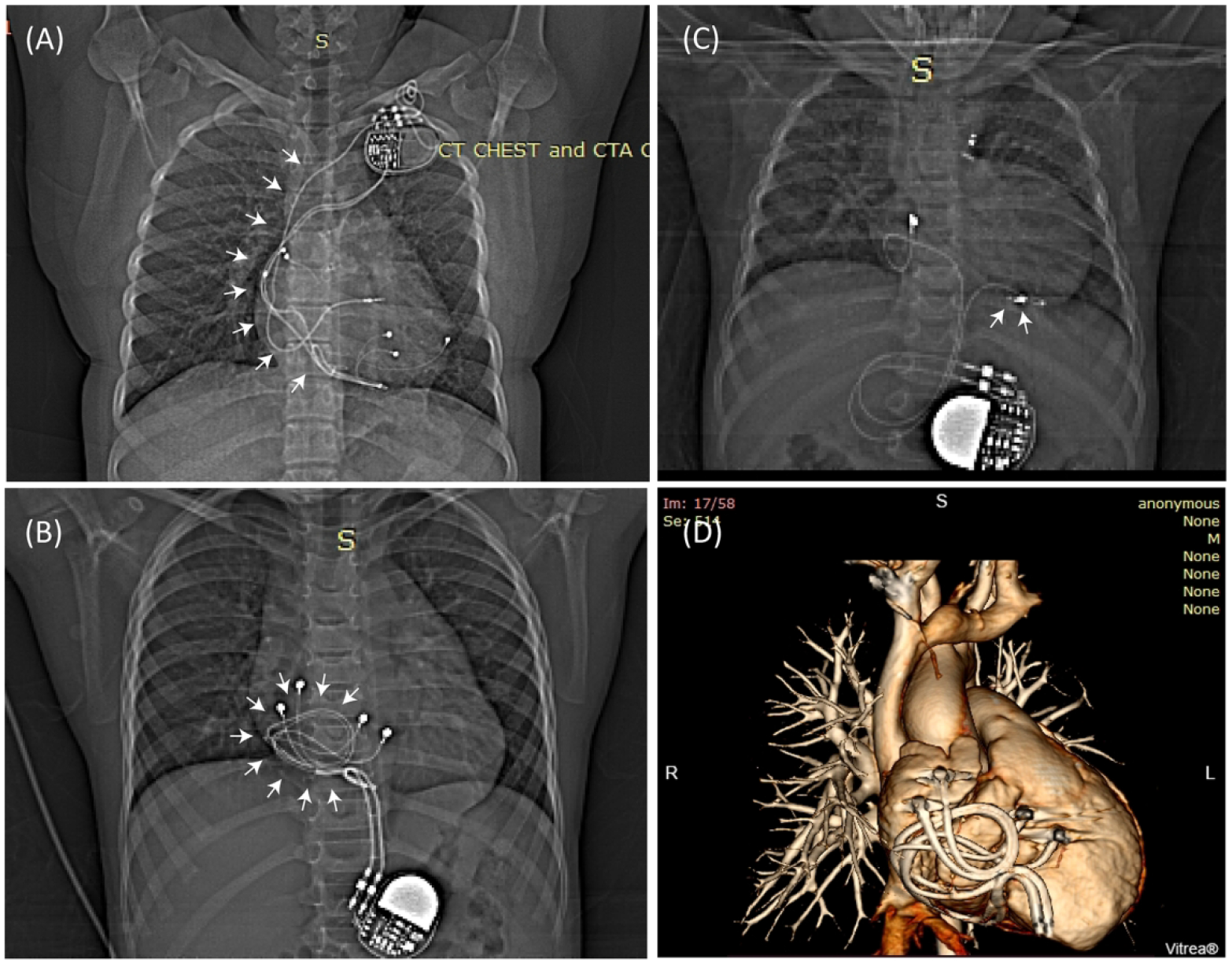
(A) Patient with an endocardial CIED. The majority of lead’s trajectory passes through the subclavian vein (white arrows) with minimal variation from patient to patient. (B) Patient with two epicardial leads looped on the anterior surface of the heart, and (C) Patient with an epicardial lead looped on the inferior surface of the heart. (D) 3D view of two epicardial leads looped on the anterior surface of the heart for (B).

In order to assess typical epicardial lead trajectories, we reviewed chest X-ray images of 100 pediatric patients implanted with CIEDs in our institution. Appropriate human subjects’ protection was provided by the respective review boards for Ann & Robert H. Lurie Children’s Hospital and Northwestern University. We identified two common categories of epicardial lead placement. In the first, excess length was looped and placed on the anterior surface of the heart (Figure 1B). In the second, the excess length of the lead was placed on or towards the inferior surface of the heart (Figure 1C). Because these two categories represented lead trajectories that were orthogonal with respect to the incident E field, we expected their RF heating to vary substantially, making one trajectory the preferred trajectory for MRI safety. Specifically, a recent study on DBS leads had shown that the variation in RF heating at lead tips could be explained by a metric that quantified the relationship between the MRI incident E field (i.e., the E field in the absence of the implant) and the lead’s orientation (16). To investigate whether such a metric could also predict RF heating of epicardial leads, we performed electromagnetic simulations to calculate the tangential component of the incident E field (referred to as E_tan_) along each representative trajectory as:

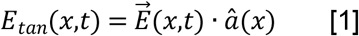

where 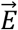 is the incident electric field and *â* is a unit vector tangential to the lead’s path (note that 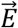 is the electric field in the absence of the implanted lead). We then calculated the peak-to-peak value of the induced voltage along each trajectory as:

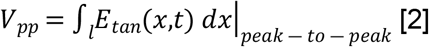

We expect that a lead placed along a trajectory with a lower *V_pp_* would generate lower SAR and lower RF heating at its tip when exposed to a B1 field.

### Abandoned leads

Over the lifetime of a CIED, leads may be disconnected and abandoned due to fracture, insulation breaks, or abnormal pacing or sensing (20,21). Although detached from the pulse generator, non-functional leads usually remain in place due to risks associated with lead extraction. Previous studies have shown an elevated risk of RF heating in abandoned leads, especially when they are capped at the proximal end (22). Theoretically, disconnecting the lead from the battery and capping (i.e., insulating) its proximal end would change the electrical length and termination boundary of the cable and thus, affecting the degree of coupling with the MRI electric field. To assess whether improvements in MRI safety due to trajectory modification would translate from fully implanted systems to abandoned leads, we also performed experiments with a capped abandoned epicardial lead.

## Method

### Electromagnetic simulations

#### Identifying epicardial lead trajectories that minimally couple with MRI incident electric field

Electromagnetic simulations were implemented in ANSYS Electronics Desktop 2020 R2 (ANSYS Inc., Canonsburg, PA). A pediatric body model consisting of average tissue, skull, brain, ribcage, and heart was created from segmented MRI images of a 29-month-old child (23) and post-processed to form a tetrahedral mesh for finite element simulations (see Figure 2). A model of a 16-rung high-pass birdcage coil with dimensions mimicking a Siemens 1.5 T Aera body coil was created based on technical specifications provided by the vendor and tuned to 63.6 MHz (24). The pediatric body model was positioned inside the coil with the heart at the magnet iso-center, and *E_tan_* and *V_pp_* were calculated along the two representative epicardial lead trajectories as shown in Figure 3. Here, green arrows show the incident electric field 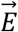 at a certain time point along each trajectory. Figure 3A also shows *E_tan_*(*t*) as a color field overlaid on each lead trajectory. Note that the magnitude of *E_tan_* is a function of time, as the orientation of the electric field changes while the field vector rotates over a full cycle. Figure 3B shows the time evolution of 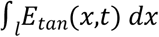 for each lead trajectory.

**Figure 2:**
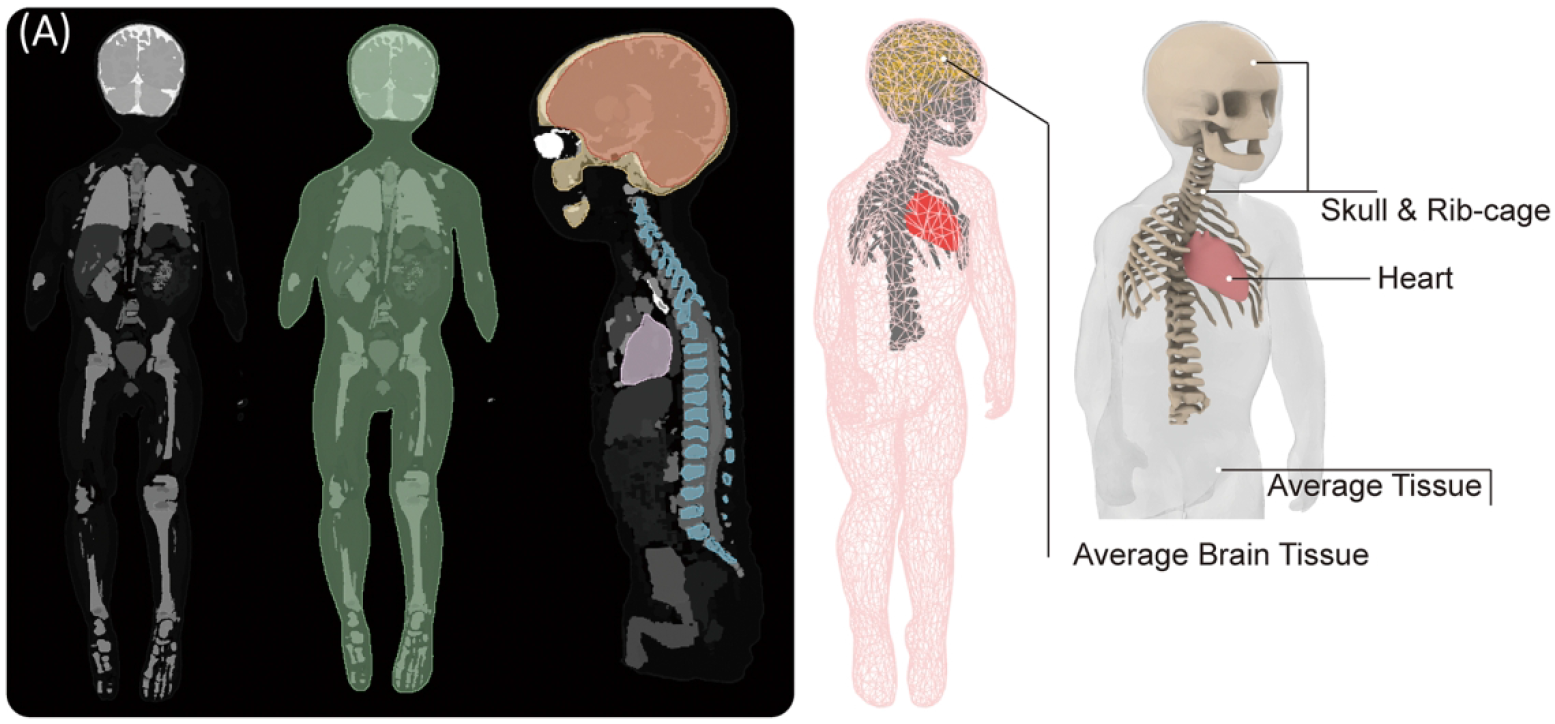
(A) Segmented MRI of a 29-month-old child was used to create 3D models of the child’s silhouette, skull, brain, ribcage, and heart. The tetrahedral mesh was post-processed for finite element simulations with four major tissue conductivities.

**Figure 3:**
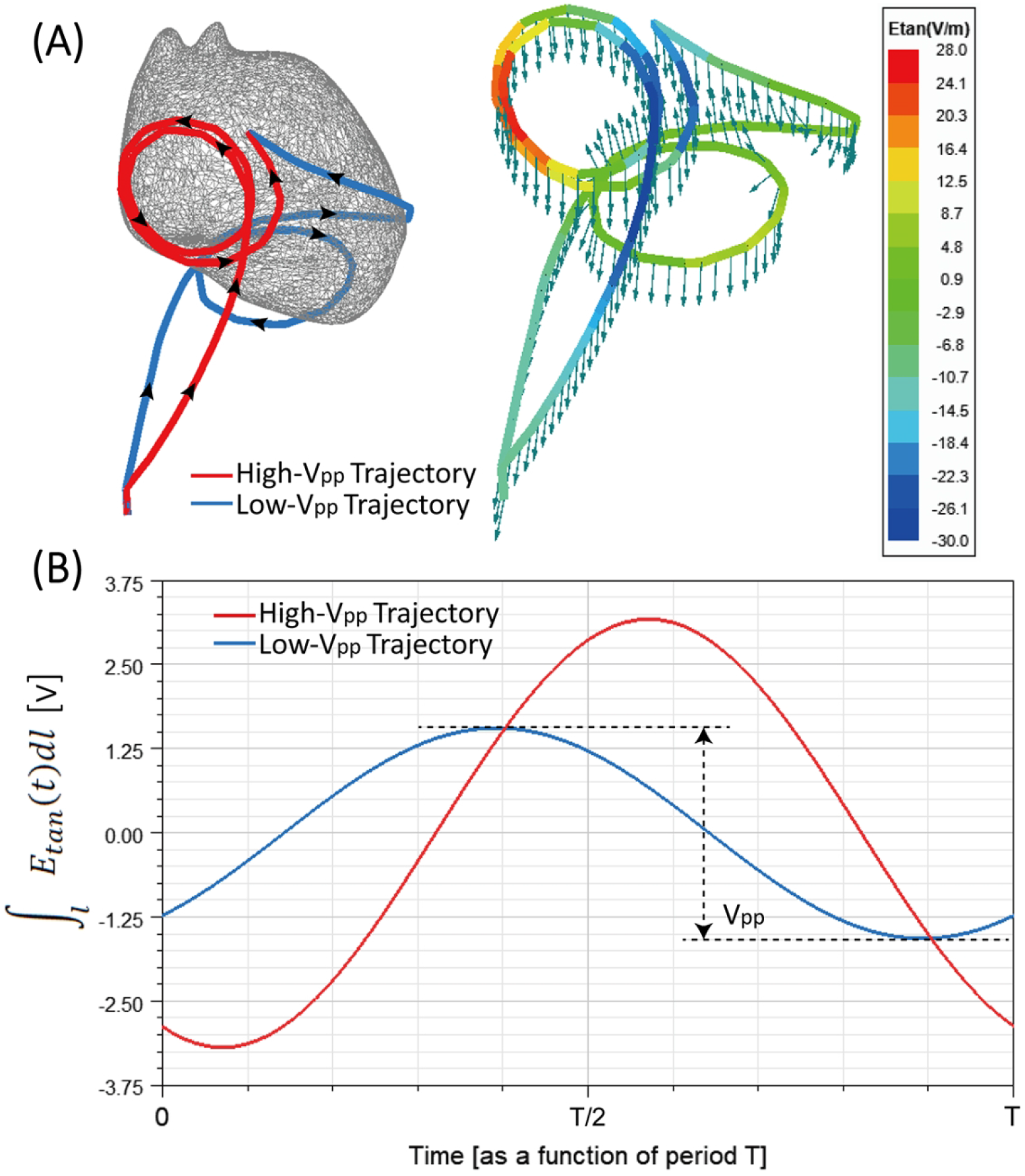
(A) Two representative epicardial lead trajectories with loops on the anterior surface (High-V_pp_ Trajectory) and inferior surface of the heart (Low-V_pp_ Trajectory) superimposed in one heart model; Green arrows show the incident electric field along the trajectory of each lead. E_tan_ is also overlaid on each lead trajectory as a color field. (B) The time evolution of induced voltage for each trajectory.

#### CIED modeling and SAR calculation

To investigate whether an epicardial lead positioned on a low-*V_pp_* trajectory deposits less power in the tissue around its tip compared to a lead placed on a high-*V_pp_* trajectory, we modeled two full CIEDs, each consisting of an implantable pulse generator (IPG) and a 35cm insulated lead, positioned on a high-*V_pp_* or low-*V_pp_* trajectory. Each lead was modeled as a solid wire made of platinum-iridium (σ= 4 × 10^6^ S/m), embedded in a 0.5-mm thick urethane insulation. The tip was modeled as a semi-spherical conductor mimicking Medtronic EPI 4965 lead model (Medtronic CapSure^®^ EPI 4965).

For both simulations, the input power of the coil was adjusted to generate an average 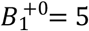 *μT* on an axial plane positioned approximately 5 mm above the center of the lead tip passing through the whole body. The 0.1g-averaged SAR (referred to as 0.1gSAR) was calculated using HFSS built-in SAR module that implements IEEE STD P1528.4 recommendation (25). The maximum of 0.1g SAR was recorded inside a cubic region of 20 mm× 20 mm× 20 mm surrounding the lead tip and was used to compare the two trajectories, as shown in Figure 4. To enhance the simulation accuracy, the initial mesh was set such that the maximum element size was < 0.5 mm on the entire lead core, < 2 mm on the IPG, in the cubical tissue region surrounding the lead tip, and the lead insulation, <10 mm in the heart, brain, ribcage, and skull tissue, and <20 mm everywhere else in the body model. ANSYS Electronics Desktop follows an adaptive mesh scheme with the successive refinement of the initial mesh between iterative passes. Scattering parameters (S-parameters) are evaluated at each port and compared to the previous pass at each adaptive pass. Simulations were considered to be converged when the magnitude of the change in S-parameters between the two consecutive passes fell below a set threshold of 0.001. All simulations converged within five adaptive passes. The convergence of absorbed RF power density was then verified by measuring the maximum of 0.1gSAR for both trajectories with different convergence thresholds (see Supporting Information Table S1), showing less than 0.3% change in the 0.1gSAR. Supporting Information Table S2 gives mesh statistics for each trajectory. Total simulation time was about 5 hours for high-*V_pp_* trajectory and 3 hours for low-*V_pp_* trajectory on a DELL server with 1.5 TB memory and 2_Xenon(R) Gold 6140 CPUs, each having 32 processing cores.

**Figure 4:**
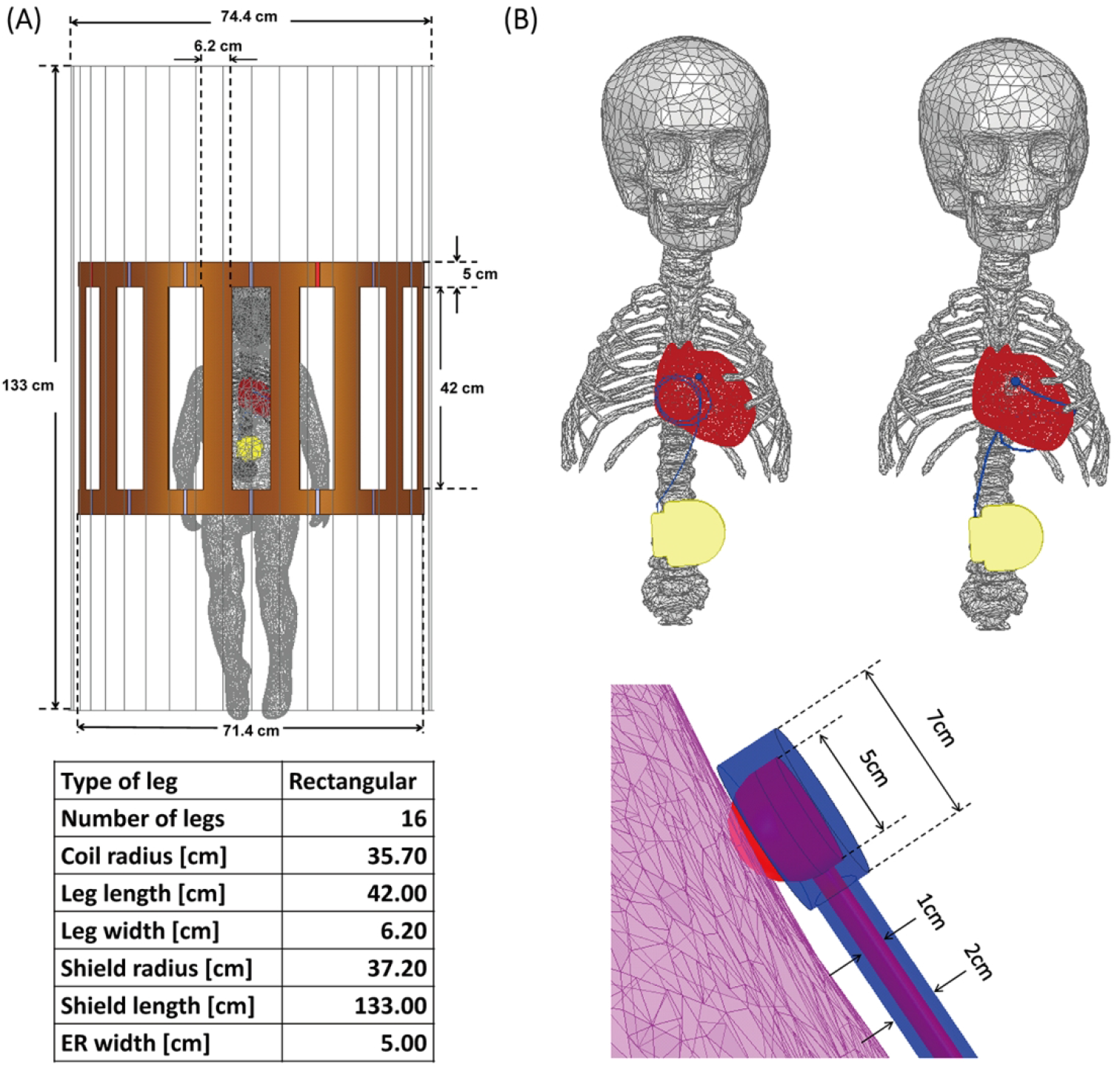
(A) The body model consisting of average tissue (σ = 0.47 S/m; ε_r_ = 80), heart (σ = 0.68 S/m; ε_r_ = 107), brain (σ = 0.40 S/m; ε_r_ = 76), skull and ribcage (σ = 0.06 S/m; ε_r_ = 16.6), placed inside a model of Siemens 1.5 T Aera body coil with the heart at the iso-center. (B) Epicardial leads with a loop positioned at the anterior and inferior surface of the heart, and a close-up of the lead tip/heart tissue interface with parameters.

### Experiments

#### Design and construction of the anthropomorphic pediatric phantom

To examine whether the results of simulations with simplified lead models would translate to experiments with realistic devices, we designed and fabricated a pediatric phantom consisting of a human-shaped container, skull, ribcage, and heart from the same MR images that were used to create the body model for simulations. Images of the 29-month-old infant were postprocessed in 3D slicer (Slicer 4.10, http://slicer.org) to generate smoothed masks of different body parts, which were further processed in a CAD tool (Rhino 6.0, Robert McNeel & Associates, Seattle, WA) to create 3D-printable objects as illustrated in Figure 5. The skull structure consisted of two separable coronal halves so that it could be filled with tissue-mimicking gel with average electrical properties of brain tissue. Similarly, a two-part mold was created from segmented images of the heart and filled with agar-doped saline solution. To create the filling solution for the heart, edible agar (Landor Trading, Montreal, Canada, gel strength = 900mg/2cm+) was gradually added to 4.0 g_NaCl_/L saline solution while the mixture was stirred on a hot plate. Once agar was fully dissolved (32g/L for the brain; 40g/L for the heart), the mixture was cooled to 60 degrees Celsius, injected into the heart mold with a syringe, and left to cool to room temperature. Once solidified, the agar-based heart structure was taken out of the mold and placed on a custom-made holder inside the phantom. The rest of the body was filled with 8 L of a gel consisting of 8g/L polyacrylamide (PAA) solution and 1.55g_NaCl_/L saline. The electric properties of tissue-mimicking material were measured using a dielectric kit (N1501A) and a vector network analyzer (Keysight Technologies, Santa Rosa, CA) to be ε_r_=76, σ=0.44 S/m for the brain tissue, ε_r_=76, σ=0.65 S/m for the heart, and ε_r_=87, σ=0.48 S/m for the PAA gel representing average tissue filling the phantom container.

**Figure 5:**
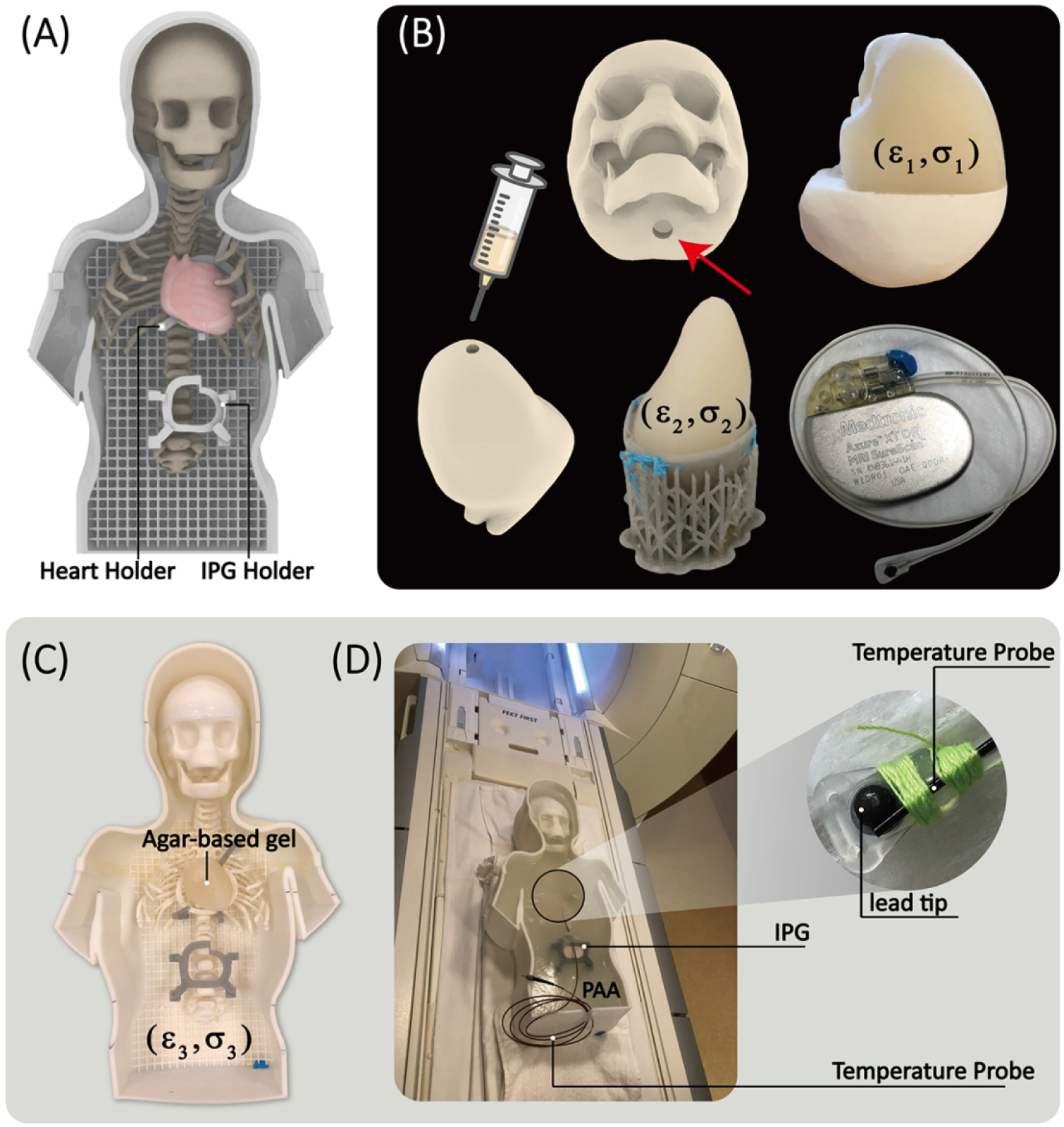
(A) Assembled phantom with two self-designed holders. (B)and(C) The skull mold was 3D printed to fill in the agar gel mimicking the brain tissue (ε_1_=76, σ_1_=0.44S/m) through the hole, enabling the conductive connection between gel inside the skull and outside. The heart mold consisted of two coronal halves attached to create a closed container filled with agar-based solution (ε_2_=76, σ_2_=0.65S/m). Once the gel solidified, the semi-solid heart structure was removed and placed on a 3D printed holder inside the phantom. The rest of the phantom container was filled with PAA mimicking average tissue (ε_3_=87, σ_3_=0.48S/m). A 35 cm epicardial lead (Medtronic CapSure^®^ EPI 4965) connected to a Medtronic Azure™ XT DR MRI SureScan pulse generator was used in the experiments. (D) Fiber optic temperature probes (OSENSA, Vancouver, BC, Canada) were secured at the tip of the lead using threads.

#### Epicardial lead trajectories and RF exposure experiments

RF heating measurements were performed in a 1.5 T Siemens Aera scanner (Siemens Healthineers, Erlangen, Germany). A 35 cm unipolar epicardial lead (Medtronic CapSure^®^ EPI 4965) was connected to a Medtronic Azure™ XT DR MRI SureScan pulse generator placed in the phantom’s abdomen approximately 10 cm caudal to the center of the heart. Fiber optic temperature probes (OSENSA, Vancouver, BC, Canada) were secured at the tip of the lead using threads. The lead’s tip was fixated to the surface of the heart phantom in a position analogous to the right atrial epicardium, and the rest of the lead was looped and placed either on the anterior surface of the heart or the inferior surface of the heart.

To assess the sensitivity of results to perturbations in lead positioning, we implemented seven slightly different instances of each representative high-*V_pp_* trajectory and low-*V_pp_* trajectory (14 trajectories in total) as illustrated in Figure 6. To examine the effect of trajectory modification on abandoned leads, we ran an additional experiment where the lead was routed on high-*V_pp_* and low-*V_pp_* trajectories, but the battery was removed, and the lead was capped at its proximal end. In all experiments, the phantom was positioned in the scanner with the heart at the iso-center, and RF heating measurements were performed using a high-SAR T1-weighted turbo spin-echo (T1-TSE) sequence (TE=7.3 ms, TR= 897 ms, B_1_^+^ = 5 μT, TA= 280 s).

**Figure 6.**
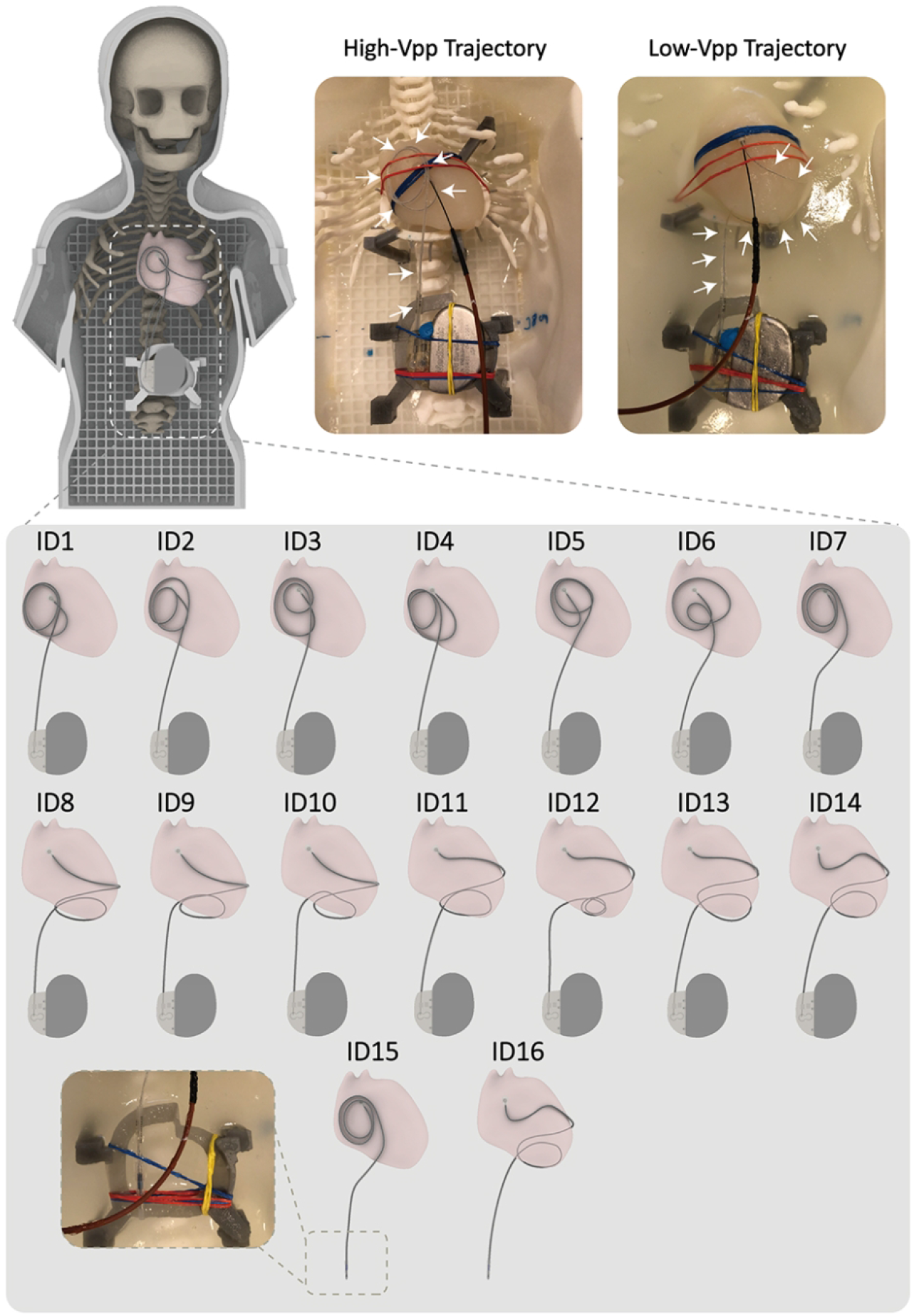
Top: The 3D printed anthropomorphic pediatric phantom and representative high-SAR (High-V_pp_ Trajectory) and low-SAR (Low-V_pp_ Trajectory) trajectories. Bottom: Each representative trajectory was replicated 7 times for sensitivity analysis. A case of abandoned lead was also examined (IDs 15&16)

## Results

### V_pp_ and SAR

For a mean B_1_^+^ = 5 μT, V_pp_ was calculated to be 6.4 volts when the loop was placed on the anterior surface of the heart and 3.1 volts for the loop on the inferior surface of the heart. The maximum of 0.1gSAR around the lead tip was recorded to be 766 W/kg for the high-*V_pp_* trajectory and 87 W/kg for the low-*V_pp_* trajectory, showing a 9-fold reduction.

### In-vitro temperature measurements: Fully implanted epicardial devices

Measured temperature rises (ΔT) at the tip of the lead for the fourteen lead trajectories are given in Supporting Information Table S3. The mean ± standard deviation of RF heating was 4.38 ± 0.43 °C for trajectories with the loop on the anterior surface of the heart and 0.27 ±0.16 °C for trajectories with the loop on the inferior surface of the heart, showing a 16-fold difference in the mean RF heating values. Figure 7 shows the temporal profile of ΔTs as well as violin plots of the data distribution. Both groups were normally distributed as calculated by the Shapiro-Wilk normality test (P>0.05). A one-tailed independent t-test at a significant level of α0 = 0.05 showed that the group with loops on the anterior surface of the heart generated significantly higher heating compared to the group with loops on the inferior surface of the heart (P-value = 9.9e-9).

**Figure 7:**
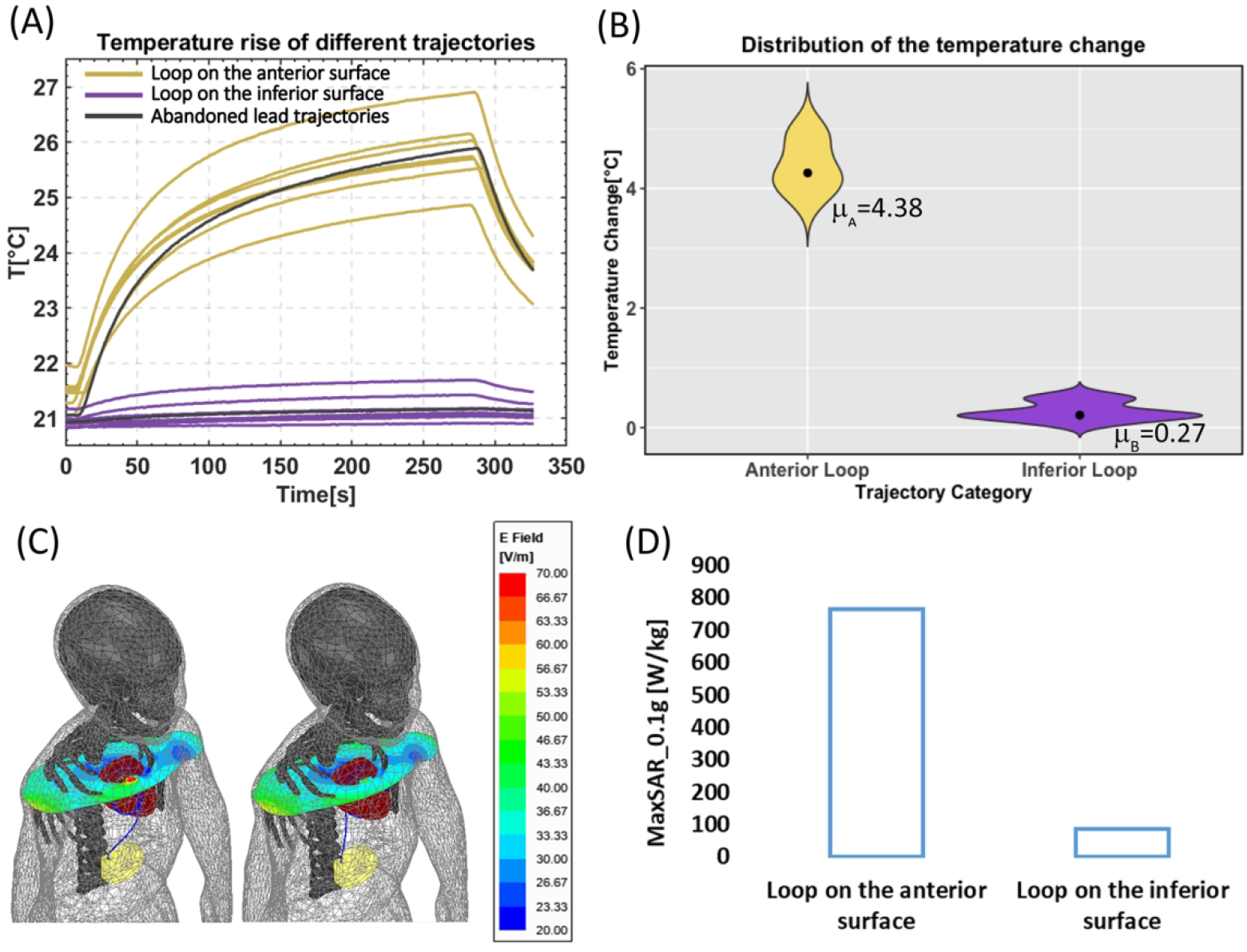
(A) Temperature rise at tips of a 35-cm epicardial lead with sixteen different trajectories shown in Figure 6. (B) Violin plots of the temperature rise distributions with the mean value for each category. (C) E field distribution on an axial plane passing through the iso-center of the coil. (D) The maximum of 0.1g-averaged SAR (MaxSAR0.1g) was calculated around the tip of the lead. The input power of the coil was adjusted to produce a mean B_1_^+^=5μT on the axial plane for both cases.

### In-vitro temperature measurements: Abandoned epicardial lead

For the abandoned capped lead, ΔT was 4.85°C for the lead routed along high-*V_pp_* trajectory and 0.24°C for lead routed along low-*V_pp_* trajectory, indicating a similar trend in the reduction of RF heating when theoretically predicted low-SAR trajectories were followed.

## Discussion

Substantial restrictions exist for patients with epicardial leads who are referred for MRI due to the elevated risk of RF heating at the tissue/lead tip interface during MRI. Recent phantom studies have reported up to 76°C of temperature rise at tips of epicardial leads during MRI at 1.5 T (3), which is concerning as children are the most likely recipients of epicardial CIED systems, and as RF leisons during early childhood can expand as the heart grows over time. (26). The problem is exacerbated by the fact that there is no straightforward method to extract epicardial leads, so children who receive these leads may be excluded from the benefits of MRI for life, even if a subsequent FDA-approved endocardial system is placed when they are older.

Efforts to make MRI accessible to patients with conductive implants have a long history, as is clear from the spate of patents published in the past two decades (27–36). Most of the work has focused on modifying the implant itself, e.g., in the case of active implants by increasing the lead’s impedance or altering the material to reduce induced RF currents (36–39). This has had limited success, and the restrictions on MRI remain. Importantly, changing the lead’s design has a substantially high cost-to-benefit ratio for the manufacturer: safety gains are modest at best, whereas slight alterations in design or materials trigger the lengthy and costly process of re-obtaining FDA approval for biocompatibility, non-toxicity, and mechanical stability. Because of this, in addition to the pediatric market is smaller than the adult market, manufacturers have been less willing to invest in MR-Conditional devices tailored for the pediatric population. Alternatively, several groups have focused on changing the MRI technology itself through field-shaping methods that create regions of the low-electric field and then steer those regions (either mechanically or electronically) to coincide with the implant (17,18,40–44). Although promising, these techniques are still in the early stages of development, and their widespread application will take years to come. Finally, recent studies have shown that vertical MRI platforms generate less heating around tips of neuromodulation devices due to orthogonal E fields and thus, could offer a safer alternative (45,46). However, studies have yet to be replicated for cardiac devices.

It is well established that the trajectory of an implanted lead and its orientation with respect to MRI electric fields substantially affects the RF heating (14,15). The idea that the lead trajectory can be manipulated at the time of implant to reduce RF heating potentially was originally suggested for neuromodulation devices (47) and later proved promising in patients with deep brain stimulation devices (16). Specifically, it was recently reported that a simple lead trajectory modification could be successfully implemented in the surgical routine of implanting deep brain stimulation leads to reduce RF heating by 3-fold during MRI at 3 T (48). Here we demonstrated that a similar simple yet highly effective technique could be adopted in children with epicardial leads to substantially reduce the risk of RF heating of the device during MRI at 1.5 T. Most importantly, we found that the reduction in RF heating was not sensitive to smaller perturbations in the lead trajectory. That is, as long as implanted leads are routed along a trajectory that roughly follows the theoretically optimum path, RF heating will be substantially reduced, even if the subtle surgical details differ from patient-to-patient. This is reassuring, as surgical constraints are likely to prevent the ideal implementation of theoretically optimal trajectories in practice. We also found that the improved safety margin would translate from fully implanted systems to abandoned leads, extending the benefits to the common clinical scenarios where lead fracture has occurred or where a patient has transitioned to an endocardial CIED, but retains the leads from the defunct epicardial system throughout life.

This study is limited in that the results cannot be extrapolated to other lead models or other MRI field strengths. This is because RF heating is a resonance phenomenon that also depends on the lead’s length, and thus, the path that minimizes the field coupling for one lead could be different for a lead with a dissimilar length. In this study, we examined the 35-cm lead because it is our pediatric hospital’s most frequently used length. However, epicardial leads come in a variety of lengths ranging from 15 cm to 85 cm, each warranting a separate investigation. In addition, we tested a unipolar lead, but some clinically implanted epicardial leads are bipolar leads which should be examined in future studies.

Overall, the results of this study indicate that a simple, easy-to-implement surgical trajectory of epicardial leads could substantially reduce their RF heating during MRI at 1.5 T in a robust and reproducible way, with benefits extending to abandoned leads.

## Acknowledgment

This work was supported, in part, by in-kind lead donations from the Medtronic Corporation (Minneapolis, MN) and by funding from the NIH (K23HL130554 and R01EB030324).

---

**Supporting Information Table S1.**

| **Loop on the anterior surface** |  |  |
| --- | --- | --- |
| **Convergence criteria(Max_ΔS)** | **No. of adaptive passes completed** | **MaxSAR0.1g(Wkg<sup>-1</sup>)** |
| 0.02 | 2 | 733.45 |
| 0.005 | 3 | 754.06 |
| 0.001 | 5 | 765.66 |
| 0.0005 | 6 | 768.09 |
| 0.0002 | 9 | 772.12 |
| **Loop on the inferior surface** |  |  |
| **Convergence criteria(Max_ΔS)** | **No. of adaptive passes completed** | **MaxSAR0.1g(Wkg<sup>-1</sup>)** |
| 0.02 | 2 | 86.66 |
| 0.005 | 3 | 86.62 |
| 0.001 | 5 | 87.06 |
| 0.0005 | 7 | 87.31 |
| 0.0002 | 8 | 87.36 |

**Supporting Information Table S2.**

| Loop on the anterior surface |  |  |  |  |
| --- | --- | --- | --- | --- |
| Parts | No. of tetrahedrons | Min. edge length(mm) | Max. edge length(mm) | RMS edge length(mm) |
| Lead | 132942 | 0.14 | 1.58 | 0.43 |
| Insulation | 222564 | 0.13 | 2.00 | 0.58 |
| IPG | 41888 | 0.32 | 7.87 | 2.35 |
| Body | 265315 | 0.14 | 27.95 | 10.28 |
| Brain | 53090 | 1.40 | 10.81 | 7.87 |
| Heart | 28758 | 0.51 | 13.25 | 6.15 |
| Rib | 29155 | 0.11 | 10.51 | 6.58 |
| Skull | 23939 | 0.42 | 10.79 | 7.62 |
| Loop on the inferior surface |  |  |  |  |
| Parts | No. of tetrahedrons | Min. edge length(mm) | Max. edge length(mm) | RMS edge length(mm) |
| Lead | 111354 | 0.11 | 1.78 | 0.44 |
| Insulation | 182920 | 0.17 | 1.99 | 0.61 |
| IPG | 41478 | 0.23 | 7.84 | 2.36 |
| Body | 259048 | 0.04 | 29.17 | 10.15 |
| Brain | 52858 | 1.13 | 12.34 | 7.86 |
| Heart | 28833 | 0.88 | 14.00 | 5.95 |
| Rib | 28106 | 0.09 | 11.10 | 6.58 |
| Skull | 23380 | 0.57 | 10.61 | 7.68 |

**Supporting Information Table S3.**

| **Trajectory ID**<br>**(from Figure 6)** | **$\Delta T$ [°C]** | **Trajectory ID**<br>**(from Figure 6)** | **$\Delta T$ [°C]** |
| --- | --- | --- | --- |
| **ID1** | **4.07** | **ID8** | **0.53** |
| **ID2** | **4.13** | **ID9** | **0.45** |
| **ID3** | **4.51** | **ID10** | **0.07** |
| **ID4** | **4.26** | **ID11** | **0.21** |
| **ID5** | **3.82** | **ID12** | **0.18** |
| **ID6** | **4.88** | **ID13** | **0.27** |
| **ID7** | **4.98** | **ID14** | **0.2** |
| **ID15**<br>**(abandoned lead)** | **4.85** | **ID16**<br>**(abandoned lead)** | **0.24** |

